# Hippocampal transcriptomic profiling reveals interferon, unfolded protein response, and synaptic signatures in RORγt-transgenic mice

**DOI:** 10.64898/2026.09.13.751197

**Authors:** Tetsuya Sasaki, Sae Sanaka, Kenyu Nakamura, Suguru Iwata, Momo Morikawa, Yosuke Takei

**Author notes:** Corresponding authors: Tetsuya Sasaki, PhD and Yosuke Takei, MD, PhD Laboratory of Anatomy and Neuroscience, Department of Biomedical Sciences, Institute of Medicine, University of Tsukuba, 1-1-1 Tennodai, Tsukuba 305-8577, Ibaraki, Japan.

## Abstract

**Aim:** RORγt-transgenic mice provide a model of sustained T helper 17-cell bias, but the associated hippocampal transcriptional profile is incompletely characterized. We investigated gene-level and gene-set expression differences in this model.

**Methods:** Bilateral hippocampal samples from 10-week-old male RORγt-transgenic mice and wild-type littermates (n = 3 per genotype) underwent bulk RNA sequencing. Gene-level differential expression was evaluated using the Empirical Analysis of DGE tool in CLC Genomics Workbench. Preranked gene set enrichment analysis examined coordinated expression differences after expression filtering and exclusion of Rorc (13,813 genes).

**Results:** Sixteen genes met a false discovery rate < 0.05 (nine higher and seven lower in transgenic mice), including *Rorc, C4b, Xbp1, Nptx2* and *Mt2*. Of 212 candidates selected using unadjusted P < 0.05 and a twofold-change threshold, 155 (73.1%) had a mean expression below 0.5 reads per kilobase of transcript per million mapped reads. Hallmark interferon-α-response, interferon-γ-response, and unfolded-protein-response gene sets showed positive enrichment. Thirteen Gene Ontology biological-process sets met a false discovery rate < 0.05, including negatively enriched sets related to excitatory postsynaptic potential, glutamatergic transmission, and calcineurin-mediated signaling. *Il17a* and *Il17f* had zero reported counts, and none of the displayed leukocyte or central nervous system cell-marker genes met false discovery rate < 0.05.

**Conclusion:** This exploratory dataset identifies interferon-response, unfolded-protein-response and synaptic transcriptional signatures associated with the RORγt-transgenic genotype. The cellular sources, causal mediators and functional consequences remain unresolved.

## INTRODUCTION

T helper 17 (Th17) cells are a CD4-positive T-cell subset whose differentiation is directed by the nuclear receptor retinoic acid-related orphan receptor γt (RORγt), encoded by *Rorc*, and whose signature cytokine is interleukin-17A (IL-17A).^1^ Beyond their established immune functions, the RORγt–IL-17A axis and its interactions with glia have been implicated in brain development and behavior.^2,3^ Maternal IL-17A contributes to autism-like phenotypes in offspring,^4^ whereas neuronal IL-17A signaling can promote sociability in models of neurodevelopmental disorders.^5^ Meningeal γδ T cells modulate anxiety-like behavior through neuronal IL-17A signaling,^6^ adoptive transfer of Th17 cells promotes depression-like behavior,^7^ and a gut-initiated Th17 response induced by dietary salt is associated with neurovascular and cognitive dysfunction.^8^ Functional IL-17 receptor subunit A in central nervous system (CNS) glia provides a potential route for immune– brain communication.^9^

Clinical studies have reported peripheral Th17/IL-17-related alterations in subsets of patients with psychiatric disorders, including increased circulating Th17 cells^10^ and peripheral *RORC* expression^11^ in drug-naïve schizophrenia. In a secondary analysis of the Combining Medications to Enhance Depression Outcomes (CO-MED) trial, pretreatment IL-17 was associated with antidepressant response in a regimen-dependent manner.^12^ These associations do not establish brain-specific mechanisms or treatment efficacy.

Experimental findings indicate context-dependent effects on the hippocampus. Meningeal γδ T-cell-derived IL-17 supported hippocampal synaptic plasticity and short-term memory under noninflammatory conditions,^13^ whereas intestinal γδ T17 cells and IL-17A were implicated in impaired hippocampal mitophagy and depression-like behavior after chronic restraint stress.^14^ These differing contexts do not imply uniformly detrimental IL-17 effects. Recent neural-circuit studies further indicate that IL-17-family effects depend on ligand identity, receptor composition and anatomical context.^15–17^

To examine brain tissue in a persistently Th17-biased background, RORγt-transgenic (Tg) mice carrying full-length murine RORγt cDNA under human CD2 regulatory elements, Tg(CD2-*Rorc*)#Staka, provide an informative model. These mice exhibit preferential transgene expression in T cells, increased IL-17-producing CD4-positive T cells, hyperglobulinemia, and autoantibody production.^18^ Circulating IL-17A was elevated at 10 and 16 weeks in previous studies on a C57BL/6 background.^19,20^ However, this constitutive model does not isolate circulating IL-17A from other systemic or developmental effects.

Previous CNS studies in this line detected relatively subtle phenotypes. At 16 weeks, dentate-gyrus Iba1 immunoreactivity and microglial density were reduced, without significant differences in astrocyte markers, doublecortin-positive immature neurons, hippocampal NR2A, NR2B, PSD-93 or PSD-95 protein levels, or novel object location performance.^20^ Behavioral work at 81–100 days reported lower body weight and altered locomotor timing, without detectable differences in aggression or conventional light–dark box measures.^21^ In pregnant Tg mice, enhanced polyinosinic–polycytidylic acid [poly(I:C)]-induced fetal loss without a corresponding acute IL-17A rise suggested that additional mediators might contribute.^19^

However, transcriptome-wide changes in untreated adult hippocampal tissue from this line remain incompletely characterized, leaving unresolved whether the genotype is associated with coordinated immune, cellular-stress or neuronal transcriptional programs. We therefore performed bulk RNA sequencing of bilateral hippocampi from untreated, 10-week-old male RORγt Tg mice and wild-type (WT) littermates. We examined gene-level differential expression, expression-filtered preranked gene set enrichment analysis (GSEA), and selected immune- and CNS-cell-marker transcripts to identify transcriptional associations, not to establish a cell-specific mechanism or causal link to behavior.

## MATERIALS AND METHODS

### Animals and hippocampal sampling

RORγt-overexpressing transgenic mice, Tg(CD2-Rorc)#Staka (MGI:5805451), were generated as described previously,^18^ using a full-length murine RORγt cDNA under human CD2 regulatory elements. The line was maintained as heterozygotes on a C57BL/6 background by backcrossing. Male Tg mice and WT littermates were studied at 10 weeks of age (n = 3 per genotype). Animals were housed under specific-pathogen-free conditions at the Laboratory Animal Resource Center, University of Tsukuba, on a 12:12-h light–dark cycle with food and water ad libitum. They received no experimental treatment before tissue collection. Mice were euthanized by cervical dislocation. Hippocampi from both hemispheres were dissected, immersed in RNAlater Stabilization Solution (Invitrogen/Thermo Fisher Scientific, AM7020), and stored at −80 °C until processing for RNA extraction. Each of the six RNA-seq samples represented one mouse.

All animal procedures were conducted in accordance with the University of Tsukuba animal-care guidelines and the National Institutes of Health Guide for the Care and Use of Laboratory Animals, under institutional animal-experiment and recombinant-DNA oversight in force at the time of the work.

WT1–WT3 correspond to analysis IDs TKS_001_01–03 and animal IDs 402, 403 and 408, respectively; Tg1–Tg3 correspond to TKS_001_04–06 and animal IDs 434, 435 and 436. Table S1 retains the original QC identifiers, which use the prefix TSK rather than TKS, together with preparation labels and sequencing run IDs. These identifier fields are preserved separately because the original QC-preparation and run-index labels are not identical for samples 05 and 06.

### RNA extraction, library preparation and sequencing

Total RNA was extracted using TRIzol reagent (Thermo Fisher Scientific/Invitrogen, 15596026) and resuspended in water. RNA was assessed using a NanoDrop spectrophotometer and an Agilent Bioanalyzer with the RNA 6000 Pico Kit (Agilent Technologies, 5067-1513). RNA integrity number (RIN) values were 8.6, 8.1 and 8.1 for WT samples and 8.2, 8.2 and 8.4 for Tg samples (range, 8.1–8.6; mean, 8.27 in each genotype; Table S1).

Library preparation and sequencing were performed by Tsukuba i-Laboratory, Transborder Medical Research Center, University of Tsukuba. Ribosomal RNA was depleted from 500 ng total RNA per sample using the NEBNext rRNA Depletion Kit (Human/Mouse/Rat; New England Biolabs, E6310), followed by directional library synthesis with the NEBNext Ultra Directional RNA Library Prep Kit for Illumina (E7420). The facility workflow specified library quality assessment using an Agilent Bioanalyzer High Sensitivity DNA Kit (5067-4626). Libraries were sequenced on an Illumina NextSeq 500 with a High Output Kit v2 using 2 × 36-bp paired-end reads (Figure 1A).

**Figure 1.**
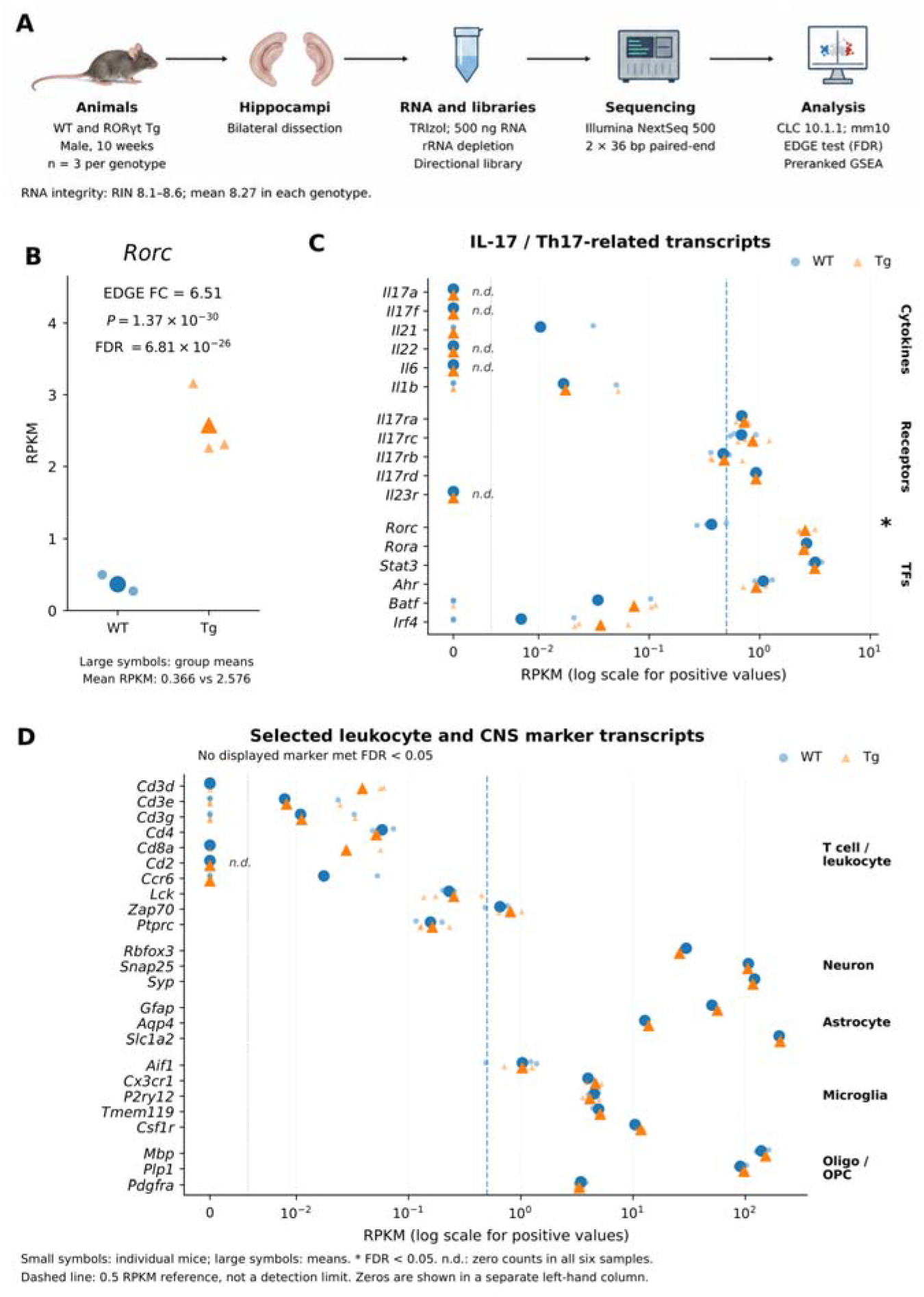
Rorc expression and selected immune- and CNS-cell-marker transcripts in WT and RORγt-transgenic mouse hippocampi. (A) Experimental workflow. Bilateral hippocampal samples were collected from 10-week-old male WT littermates and RORγt-transgenic (Tg) mice (n = 3 per genotype). After cervical dislocation, hippocampi from both hemispheres were dissected, immersed in RNAlater (AM7020), and stored at −80 °C until processing. RNA was extracted with TRIzol; RIN values ranged from 8.1 to 8.6 (mean, 8.27 per genotype). Directional libraries were prepared from 500 ng total RNA after rRNA depletion and sequenced on an Illumina NextSeq 500 using 2 × 36-bp paired-end reads. Reads were analyzed in CLC Genomics Workbench v10.1.1 against mm10, followed by gene-level differential expression analysis and preranked GSEA. (B) Individual-animal Rorc RPKM values. The fold change shown (6.51, Tg/WT) is the EDGE estimate; the ratio of the group mean RPKM values (0.366 and 2.576) is 7.03. (C) Selected IL-17/Th17-related cytokines, receptors and transcription factors. Il23r is classified as a receptor. Il21 had a count of 1 in WT1 (RPKM = 0.0312) and zero counts in the other five samples. (D) Selected T-cell/leukocyte and CNS-cell-marker transcripts; no displayed gene met FDR < 0.05. In (B–D), circles denote WT and triangles denote Tg; small symbols represent individual mice and large symbols show arithmetic group means including zeros. Positive RPKM values in (C,D) are shown on a logarithmic scale; zeros occupy a separate column labeled 0, without a pseudocount. n.d. denotes zero reported counts in all six samples. The 0.5-RPKM dashed line is an expression-filter reference, not an experimentally established detection limit. Statistical values are taken from the original EDGE output; FDR was not recalculated within the displayed subset. *FDR < 0.05. These data do not establish equivalent cell proportions or exclude rare-cell infiltration or low-abundance cytokine expression. Abbreviations: bp, base pairs; CNS, central nervous system; FC, fold change; FDR, false discovery rate; GSEA, gene set enrichment analysis; IL, interleukin; OPC, oligodendrocyte precursor cell; oligo, oligodendrocyte; RIN, RNA integrity number; RNA-seq, RNA sequencing; rRNA, ribosomal RNA; RPKM, reads per kilobase of transcript per million mapped reads; TFs, transcription factors; WT, wild type. EDGE denotes the Empirical Analysis of DGE tool in CLC Genomics Workbench. Panel A is an illustrative workflow schematic; quantitative panels B–D are generated from the source expression data.

### Read processing, alignment, and gene-level differential expression

According to the facility workflow, FASTQ files from the imaging sections were concatenated separately for Read 1 and Read 2 and imported as paired-end reads. Reads were mapped to mm10 using the Ensembl-derived gene and transcript annotation distributed with CLC Genomics Workbench v10.1.1 (QIAGEN). The gene-level output contained 49,585 annotated features; the workflow listed 103,982 transcript isoforms. Sample-level read and mapping counts are provided in Table S1. Mapped-read totals are the sum of reads reported as mapped in pairs and mapped in broken pairs; all these quantities are reads, not read pairs.

Genotype differences were assessed using the Empirical Analysis of DGE (EDGE) tool in CLC, described in the original analysis as an implementation of the edgeR exact test with tagwise dispersions.^22,23^ Gene-level P values and Benjamini–Hochberg false discovery rate (FDR)-adjusted P values were retained from the supplied analysis output and were not recalculated within displayed or expression-filtered subsets. FDR < 0.05 defined the primary gene-level criterion, without an additional fold-change requirement. For a separate exploratory comparison, candidates were selected using unadjusted P < 0.05 and a change of at least twofold in either direction.

CLC encodes a decrease as a negative inverse fold-change ratio. Accordingly, log□ fold change was calculated as sign(FC_CLC) × log□(|FC_CLC|), where FC_CLC denotes this signed ratio. The exploratory twofold criterion is therefore |FC_CLC| ≥ 2, equivalently |log□ fold change| ≥ 1. Reads per kilobase of transcript per million mapped reads (RPKM) were used for abundance summaries and visualization; they were not substituted for total counts in the original count-based test. Group means include zero values. The facility supplied quantile-normalized diagnostic plots separately; the principal component analysis (PCA), correlations and gene-expression heatmaps reported here used the exported RPKM values rather than the quantile-normalized columns (Table S2). Gene-level summaries and expression displays are presented in Figure 2.

**Figure 2.**
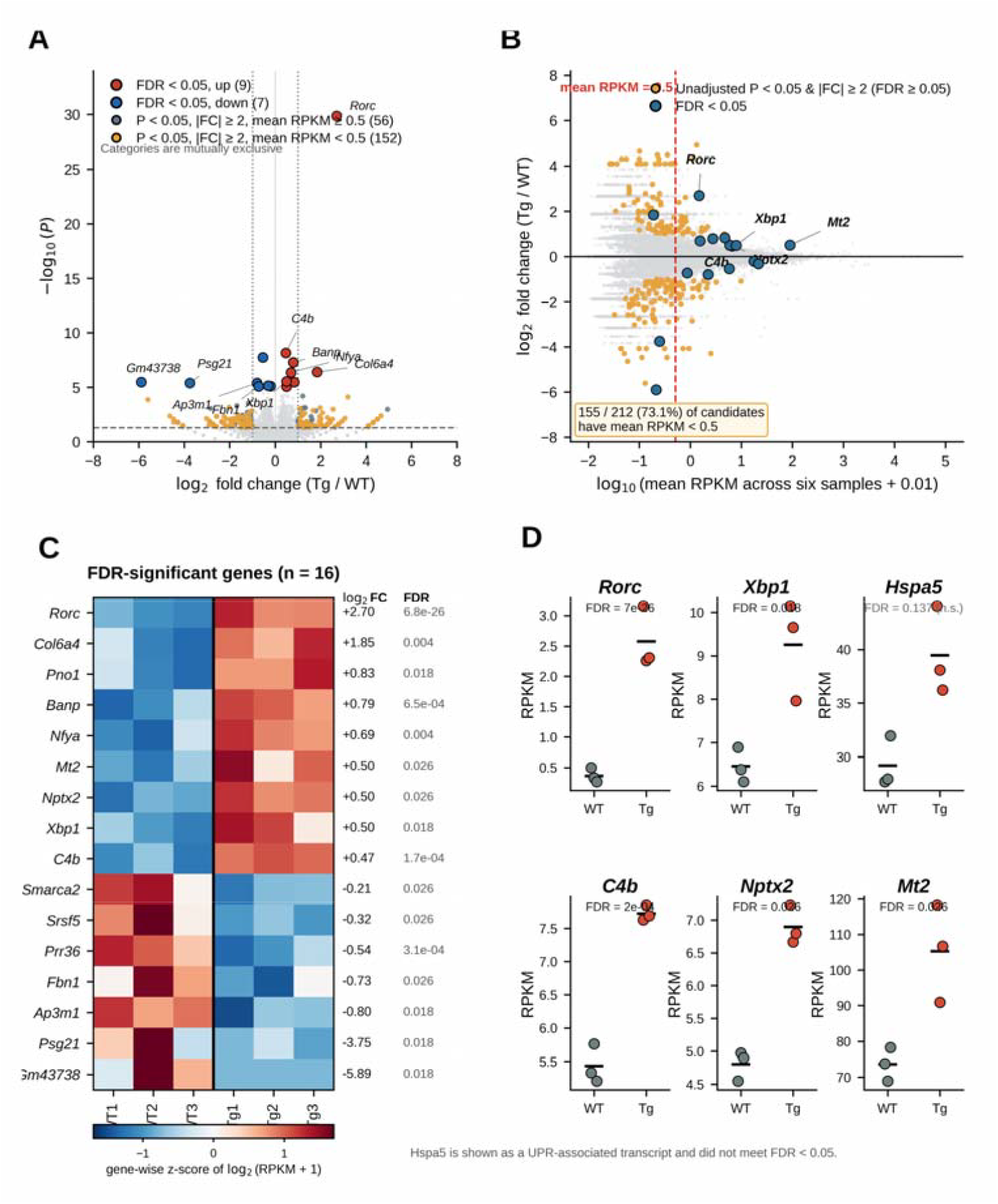
Gene-level differential expression and the abundance of candidates selected using unadjusted P values and fold change. (A) Volcano plot of the original EDGE results. Nine genes showed higher and seven showed lower expression in Tg samples at FDR < 0.05, without a fold-change requirement. Other candidates meeting unadjusted P < 0.05 and |CLC-signed fold change| ≥ 2 are colored by mean RPKM across all six samples. Categories are mutually exclusive: genes meeting the FDR criterion are assigned first to the FDR-significant categories. The horizontal dashed line indicates unadjusted P = 0.05; vertical dotted lines indicate |log□ fold change| = 1. (B) Mean expression across all six samples versus EDGE log□ fold change. The vertical dashed line indicates mean RPKM = 0.5. Of 212 unadjusted-P/twofold candidates, 155 (73.1%) had a mean RPKM < 0.5. FDR-significant genes are distinguished from the other candidates. (C) Gene-wise z-scores of log□(RPKM + 1) for all 16 FDR-significant genes, with EDGE log□ fold change and FDR at right. (D) Individual-animal RPKM values for selected genes; horizontal bars show group means. Hspa5 is included as a UPR-associated transcript that did not reach FDR < 0.05 (FDR = 0.137). WT and Tg each comprise three individual mice. All inferential statistics are retained from the original analysis. Abbreviations: FC, fold change; FDR, false discovery rate; n.s., not significant at FDR < 0.05; RPKM, reads per kilobase of transcript per million mapped reads; UPR, unfolded protein response; WT, wild type; Tg, RORγt transgenic. EDGE denotes the Empirical Analysis of DGE tool in CLC Genomics Workbench.

### Principal component analysis and sample correlations

Sample-level quality control (QC) retained genes with a mean RPKM > 0.5 across all six samples (13,415 genes). After log□(RPKM + 1) transformation, the 2,000 genes with the highest variance were selected for PCA. Each gene was centered across samples without scaling to unit variance. Pearson correlations were calculated using all 13,415 filtered genes, not only the 2,000 PCA genes. All six samples were retained, and Rorc was not excluded from the QC-filtered gene set, in contrast to its exclusion from GSEA. No inferential comparison of within-genotype and between-genotype correlations was performed (Figure S1).

### Expression filtering and gene ranking

For preranked GSEA, genes were retained if their mean RPKM exceeded 0.5 in at least one genotype, yielding 13,814 genes. Genes were ranked in descending order using sign(log□ fold change) × −log□□(P), with unadjusted P values from the EDGE analysis and positive scores indicating higher expression in Tg samples. This GSEA expression filter was distinct from the criterion used for sample-level QC and was not applied retrospectively to redefine the 16 FDR-significant genes. The gene-level *Rorc* entry was excluded before enrichment analysis, leaving 13,813 genes in the input ranking. The complete expression-filtered ranking is provided in the Ranked_gene_list worksheet of Table S3, with used_in_GSEA set to TRUE for the 13,813 included genes and FALSE for *Rorc*. The GSEA_input_noRorc worksheet contains only the included genes and their ranking scores.

### Preranked gene set enrichment analysis

Preranked GSEA^24^ was performed using GSEApy v1.3.1^25^ and Molecular Signatures Database (MSigDB) mouse collections v2025.1.Mm.^26^ The archived outputs comprise 50 Hallmark sets, 3,595 Gene Ontology biological-process (GO-BP) sets and 1,553 mixed curated sets. The last collection includes pathway definitions and experimental gene signatures; it is not treated as a Reactome/WikiPathways-only collection. Gene sets were tested against the 13,813-gene input ranking; eligible sets contained 15–500 members represented in that ranking. Available analysis records document 1,000 gene-set permutations. The exact original GMT filenames, random seed, weighting parameter and execution-environment details were not retained in the archived materials available for this preprint.

Enrichment was summarized by the normalized enrichment score (NES), nominal P value and FDR q value in the original testing context of each collection. Positive NES denotes enrichment toward the Tg-associated end of the ranked list, and negative NES denotes enrichment toward the opposite end. FDR q < 0.05 was used for the primary gene-set interpretation. Exported zero nominal P values were retained as permutation estimates and were not interpreted as exact zero probabilities. No FDR recalculation was performed after selecting subsets for figures or grouping terms by name. Complete exported results are supplied in Table S3.

Figure 3A displays representative GO-BP terms with nominal P < 0.01 and at least five leading-edge genes; all 13 GO-BP terms meeting FDR q < 0.05 are shown in Figure S2. Figure 3B displays selected Hallmark terms with nominal P < 0.10 and distinguishes FDR-significant from non-significant sets. Figure 3D displays the first ten genes in the exported leading-edge list for each of the four sets illustrated in Figure 3C, retaining that order. This correspondence was verified against the complete result table; it is not a selection of the ten largest expression effects. The 40 displayed columns represent 39 unique genes because *Gsk3b* appears in two blocks. Each gene was standardized across the six samples after log□(RPKM + 1) transformation, using its population standard deviation (ddof = 0). Complete leading-edge memberships are supplied in Tables S3 and S4.

**Figure 3.**
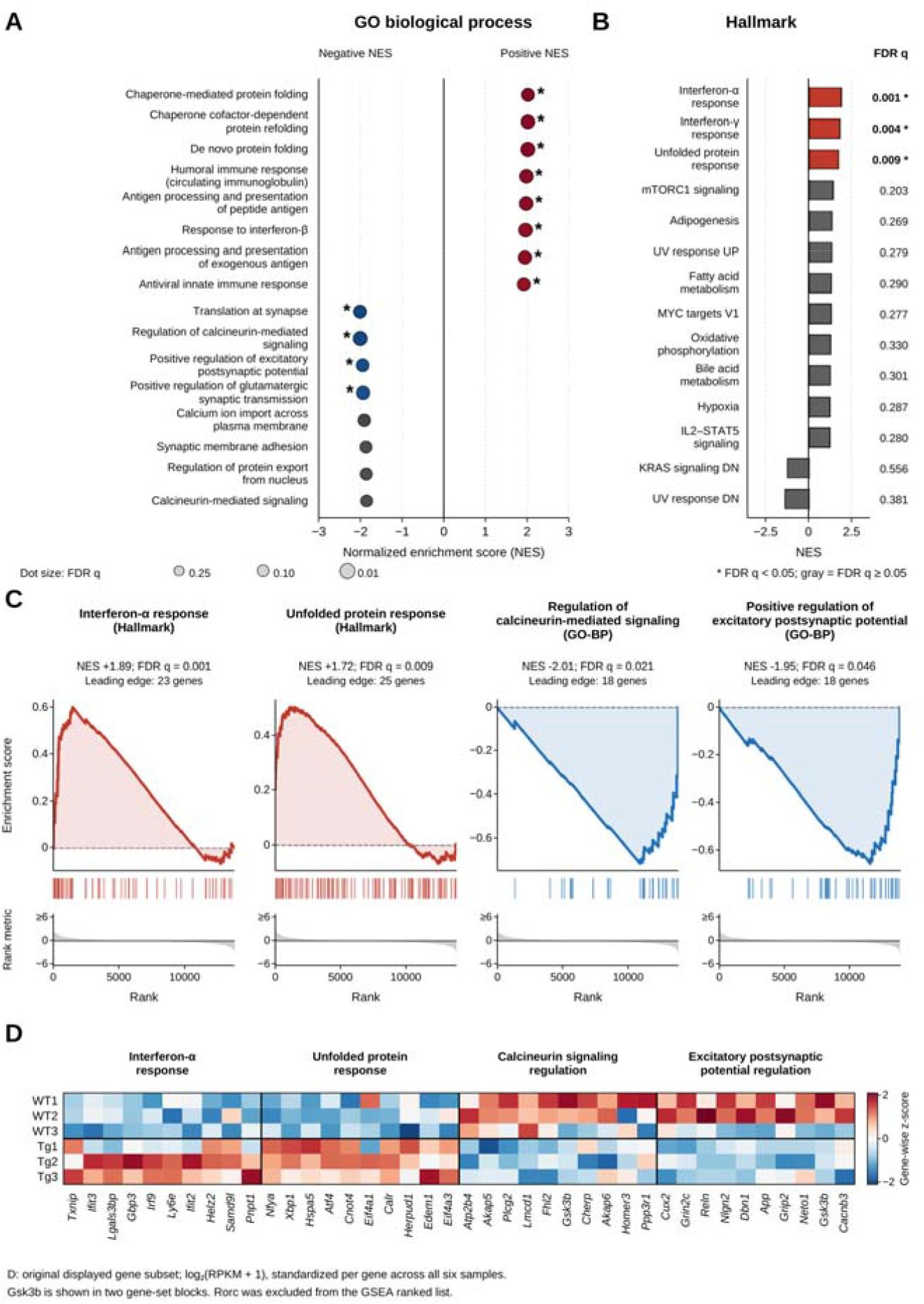
Interferon-response, unfolded-protein-response and synaptic gene-set enrichment in RORγt-transgenic mouse hippocampi. (A) Representative GO biological-process terms from the preranked GSEA, selected among terms with nominal P < 0.01 and at least five leading-edge genes. Dot position indicates NES, dot size reflects FDR q, and asterisks denote q < 0.05. Non-significant terms are gray. All 13 FDR-significant GO-BP terms are shown in Figure S2 and Table S3; the present panel is not an exhaustive list. (B) Selected Hallmark sets with nominal P < 0.10. Bars indicate NES and the adjacent column shows rounded FDR q values. The three positive enrichments meeting q < 0.05 are colored; other sets are gray. Positive NES denotes enrichment toward the Tg-associated end of the ranking, and negative NES denotes enrichment toward the opposite end. (C) Running enrichment scores for the Hallmark IFN-α response and unfolded protein response sets, and the GO sets regulation of calcineurin-mediated signaling and positive regulation of excitatory postsynaptic potential. Vertical ticks indicate ranked gene-set members; lower tracks show the ranking metric. The reported NES, q values and leading-edge counts are retained from the original analysis. (D) The first ten genes in each corresponding exported leading-edge list, retained in exported order. This displays 40 columns representing 39 unique genes because Gsk3b occurs in two blocks. Colors are gene-wise z-scores of log□(RPKM + 1) across WT1– WT3 and Tg1–Tg3, calculated with population standard deviations (ddof = 0). The color range is −2.1 to +2.1; no displayed value is clipped. The heatmap summarizes the same analyzed animals, not independent validation. Full-precision statistics and complete leading edges are provided in Tables S3 and S4. Rorc was excluded before GSEA, leaving 13,813 genes in the input ranking. The complete pre-exclusion ranking and used_in_GSEA flags are provided in Table S3. Abbreviations: FDR, false discovery rate; GO-BP, Gene Ontology biological process; GSEA, gene set enrichment analysis; IFN, interferon; LE, leading edge; NES, normalized enrichment score; RPKM, reads per kilobase of transcript per million mapped reads; WT, wild type; Tg, RORγt transgenic.

### Leading-edge overlap

To summarize redundancy among significant enrichments, leading-edge overlap was calculated for the three FDR-significant Hallmark sets and 13 FDR-significant GO-BP sets. For two leading-edge sets A and B, Jaccard similarity was |A ∩ B|/|A ∪ B|. The matrix was ordered by average-linkage hierarchical clustering on 1 − Jaccard, followed by optimal leaf ordering. This analysis describes shared leading-edge membership rather than GO semantic similarity or an additional significance test (Figure S3; Table S4).

### Software and analytical scope

The CLC gene-level statistics and the original GSEA enrichment statistics were retained during manuscript and figure revision. Subsequent tabulation, abundance summaries, PCA, sample correlations and heatmap calculations used Python-based scripts; the source values and derived quantities are provided with the supplementary tables and figure data. These descriptive recalculations did not constitute a new EDGE or GSEA analysis. Version-specific dependencies will accompany the public code archive when finalized.

### Use of generative artificial intelligence

Claude (Anthropic) was used to assist with analytical coding and literature information handling. ChatGPT (OpenAI) was used during subsequent data-table checking, descriptive QC calculations, literature searching, and reference formatting. The authors take full responsibility for the analysis, figures, references and text.

## RESULTS

### RNA quality and sample-level expression profiles

All six samples had RIN values between 8.1 and 8.6, with the same mean of 8.27 in the two genotypes. The mapping summary contained 26.21–30.47 million reads per sample, of which 86.01%–90.58% were reported as mapped when paired and broken-pair categories were combined (Table S1). In the PCA of the 2,000 most variable expression-filtered genes, PC1 and PC2 explained 25.3% and 20.6% of the variance, respectively (45.9% combined; Figure S1A). Pearson correlations across all 13,415 QC-filtered genes ranged from 0.9837 to 0.9875 for the 15 distinct sample pairs (Figure S1B). These summaries describe the observed expression profiles; they do not establish equivalence between genotypes or exclude gene-specific differences.

### Rorc expression and IL-17-related transcripts

*Rorc* transcript abundance was higher in Tg than in WT hippocampi (mean RPKM,0.366 versus 2.576; EDGE fold change, 6.51; P = 1.37 × 10□^3^□; FDR = 6.81 × 10□^2^□; Figure 1B). The ratio of the two group mean RPKM values was 7.03, which differs from the fold-change estimate reported by EDGE. Gene-level bulk RNA-seq does not distinguish endogenous *Rorc* isoforms from transgene-derived transcripts or identify the cells contributing to this increase. *Il17a, Il17f, Il22, Il6* and *Il23r* had zero reported counts in all six samples. In contrast, *Il21* had a count of 1 in WT1 (TKS_001_01; RPKM = 0.0312), with zero counts in the other five samples. No IL-17 receptor gene or other transcription factor displayed in Figure 1C, apart from *Rorc*, met FDR < 0.05. Mean *Il17ra* RPKM was 0.687 in WT and 0.722 in Tg (P = 0.842), and mean *Il17rc* RPKM was 0.684 and 0.864, respectively (P = 0.318). Zero counts under these sequencing and mapping conditions do not establish the absence of local cytokine production.

### Selected leukocyte and CNS-cell-marker transcripts

None of the selected leukocyte or CNS-cell-marker genes displayed in Figure 1D met FDR < 0.05. *Rbfox3* showed a nominal difference (P = 2.88 × 10□□) that was not significant after multiple-testing correction (FDR = 0.373). The gene-level results for markers of T cells/leukocytes, neurons, astrocytes, microglia, oligodendrocytes and oligodendrocyte precursors are available in Table S2. These measurements do not quantify cell proportions and cannot exclude rare-cell infiltration or modest shifts in cellular composition. Relative expression levels of *Rorc* and different lineage-marker genes were not used to infer the cellular source of the *Rorc* signal.

### Gene-level differences and the influence of low abundance

Sixteen genes met FDR < 0.05, with nine showing higher and seven showing lower expression in Tg hippocampi (Figures 2A and 2C; Table S2). In addition to *Rorc*, these included *C4b* (mean RPKM, 5.43 versus 7.72; FDR = 1.73 × 10□□), *Xbp1* (6.46 versus 9.25; FDR = 0.0185), *Nptx2* (4.81 versus 6.90; FDR = 0.0264) and *Mt2* (73.64 versus 105.28; FDR = 0.0264). *Hspa5*, displayed as an unfolded protein response (UPR)-associated transcript, had higher mean RPKM in Tg samples (29.21 versus 39.46) but did not meet the gene-level FDR criterion (FDR = 0.137; Figure 2D).

The exploratory criterion of unadjusted P < 0.05 and a change of at least twofold in either direction selected 212 genes. Of these, 155 (73.1%) had a mean RPKM < 0.5 across all six samples, and 189 (89.2%) had a mean RPKM < 1. Four of the 212 candidates—*Rorc, Col6a4, Psg21* and *Gm43738*—also met FDR < 0.05. Figure 2A uses mutually exclusive color categories, assigning these four genes to the FDR-significant categories. These results show that many candidates selected using unadjusted P values and a large fold-change threshold were low-abundance transcripts in this dataset. We therefore interpreted large fold changes cautiously and used expression-filtered gene ranking, rather than a fold-change cutoff alone, for gene-set analysis.

### Positive enrichment of interferon-response and unfolded-protein-response sets

Three of the 50 Hallmark sets met FDR q < 0.05, all with positive NES: interferon-α response (NES = 1.89, q = 0.00149; 23 leading-edge genes), interferon-γ response (NES = 1.82, q = 0.00447; 42 leading-edge genes) and the unfolded protein response (NES = 1.72, q = 0.00894; Figures 3B and 3C; 25 leading-edge genes). The interferon (IFN) leading edges included *Ifit2, Ifit3, Irf9, Isg15, Gbp3, Ly6e, Lgals3bp* and *Cd74*; *Stat1* was present in the interferon-γ leading edge. The UPR leading edge included *Nfya, Xbp1, Hspa5, Atf4, Calr, Herpud1, Edem1, Pdia6* and *Dnajc3* (Table S4). Figure 3D illustrates subsets of these genes in the same six animals and is not an independent validation cohort.

Thirteen of the 3,595 GO-BP sets met FDR q < 0.05: nine with positive and four with negative NES (Table S3; Figure S2). Positively enriched sets included response to interferon-β (NES = 1.96, q = 0.0253), antiviral innate immune response (NES = 1.92, q = 0.0371), antigen processing and presentation of peptide antigen (NES = 1.97, q = 0.0240) and chaperone-mediated protein folding (NES = 2.02, q = 0.0432). Figure 3A shows representative terms; Figure S2 includes all 13 significant sets, including cellular response to cadmium ion (NES = 1.90, q = 0.0420; nine leading-edge genes). The latter name identifies the annotated gene set and does not indicate cadmium exposure in these animals.

### Negative enrichment of synaptic and calcineurin-related gene sets

The four negatively enriched GO-BP sets meeting FDR q < 0.05 were regulation of calcineurin-mediated signaling (NES = −2.01, q = 0.0207), translation at synapse (NES = −2.01, q = 0.0404), positive regulation of glutamatergic synaptic transmission (NES = −1.94, q = 0.0422) and positive regulation of excitatory postsynaptic potential (NES = −1.95, q = 0.0459; Figures 3A and 3C; Figure S2). Synaptic membrane adhesion had a negative NES but did not meet the FDR threshold (NES = −1.87, q = 0.078).

The leading edge for positive regulation of excitatory postsynaptic potential included *Grin1, Grin2c, Neto1, Neto2, Nlgn1, Nlgn2, Nlgn3, Nrxn1, Reln, Dbn1* and *Grip2*. The calcineurin-regulation leading edge included *Akap5, Akap6, Ppp3r1* and *Homer3*. Translation at synapse was supported predominantly by cytoplasmic ribosomal protein genes (Table S4), so its enrichment cannot be assigned specifically to local synaptic translation. Membership in an enriched set does not imply that each member is individually differentially expressed. *Grin2a, Grin2b* and *Dlg4* did not meet the gene-level FDR criterion (unadjusted P = 0.510, 0.461 and 0.332, respectively; Table S2).

### Mixed curated results and overlap between gene sets

Ten of the 1,553 mixed curated sets met FDR q < 0.05, all with positive NES. These included the Reactome sets antigen presentation: folding, assembly and peptide loading of class I MHC (NES = 1.95, q = 0.0186) and cargo concentration in the ER (NES = 1.86, q = 0.0400). The collection also contained experimental signatures, including immune-checkpoint-related and other named signatures; these were not interpreted as diagnoses or direct evidence of the experimental exposures used to define them. All results, including non-significant sets, are provided in Table S3.

The IFN-α and IFN-γ Hallmark leading edges shared 21 genes among 44 genes in their union (Jaccard similarity, 0.477). Chaperone cofactor-dependent protein refolding and de novo protein folding shared seven genes among ten in their union (Jaccard similarity, 0.700). Figure S3 and Table S4 show these overlaps for all significant Hallmark and GO-BP sets. Accordingly, related enrichments were interpreted as overlapping transcriptional summaries rather than independent replications of the same mechanism.

The GO complement activation set showed a positive NES but did not meet FDR q < 0.05 (NES = 1.84, q = 0.0954). Its leading edge included *C4b, C1qa, C1qb, C1qc, Cfh* and *Cfp*. The Hallmark complement set was also not significant (NES = 1.07, q = 0.589; Table S3). Thus, the individual-gene increase in *C4b* did not establish significant enrichment across both complement collections.

## DISCUSSION

This exploratory study provides a transcriptome-wide view of the hippocampus in RORγt-transgenic mice and identifies a coherent set of genotype-associated signals. The principal findings were positive enrichment of interferon-response and unfolded protein response (UPR) gene sets, preferential WT-side enrichment of selected synaptic and calcineurin-related gene sets, and a focused group of false discovery rate (FDR)-significant genes including *Rorc, C4b, Xbp1, Nptx2* and *Mt2*. Together, these results define several convergent molecular themes that can guide mechanistic follow-up in this Th17-biased model. The strongest of these themes was the interferon-related signature.

Interferon-response enrichment was among the clearest gene-set findings. The IFN-α and IFN-γ leading edges contained overlapping interferon-responsive genes and were accompanied by positive enrichment of antigen-presentation sets. This convergence supports a shared interferon-responsive transcriptional program and strengthens the broader immune-response theme. Previous work in this line proposed contributions from mediators other than IL-17A, including interferons, to enhanced maternal poly(I:C) responses.^19^ The present data therefore identify interferon-related signaling as a priority candidate for follow-up in this line. The distinct experimental contexts—challenged pregnant females in the earlier study and untreated male hippocampal samples here— also make direct measurements of circulating interferons and downstream protein activation especially informative. This immune-response framework, in turn, provides a useful context for interpreting the detectable IL-17 receptor transcripts.

The detectable *Il17ra* and *Il17rc* transcripts provide a useful bridge between the present hippocampal dataset and recent receptor-defined neural-circuit studies. Neural IL-17-family effects have been mapped to amygdala anxiety-related circuits^15^ and cortical IL-17E-dependent social circuits,^16^ while studies of stress-related hippocampal mitophagy^14^ and protective cerebellar IL-17A effects^17^ further illustrate the importance of biological context. Taken together, these findings suggest that ligand identity, receptor composition, anatomical site and cellular source are likely to shape the consequences of IL-17-family signaling. Resolving which peripheral or local cell populations contribute in this model will therefore be an informative next step, particularly in relation to the second major transcriptional theme, the UPR.

A second prominent finding was the UPR-associated transcriptional signature, supported by *Xbp1* and multiple folding- and endoplasmic-reticulum-associated transcripts. *Xbp1* also met the individual-gene FDR threshold, whereas *Hspa5* showed a higher group mean and contributed to the UPR leading edge without reaching the gene-level FDR threshold. This combination of gene-level and gene-set evidence highlights protein-homeostasis pathways as a focused target for validation. Because gene-level *Xbp1* abundance does not distinguish spliced from unspliced forms, assays of XBP1 splicing, protein activation and additional UPR markers could directly test this transcriptional signal. Reduced body weight has been reported in this line,^21^ and IL-17 has been linked to adipogenesis and weight regulation.^27^ At the same time, mTORC1 signaling, adipogenesis and fatty acid metabolism did not meet the Hallmark FDR criterion, suggesting that the UPR-associated signal is more selective than a generalized metabolic shift. This cellular-stress context is relevant to the synaptic transcriptional pattern observed in parallel.

The synaptic and calcineurin-related gene sets add a neuronal dimension to the immune and cellular-stress signatures. Negative normalized enrichment scores indicate preferential representation toward the WT-associated end of the ranking rather than uniform downregulation of every member, and the implicated sets contained adhesion, receptor-associated and scaffolding genes relevant to psychiatric disorders.^28^ The lack of significant *Grin2a, Grin2b* and *Dlg4* differences is compatible with earlier protein measurements,^20^ whereas the increase in *Nptx2* provides a complementary signal. NPTX2 participates in AMPA-receptor clustering and excitatory synapse maturation^29,30^ and can restrain complement-associated synapse loss.^31^ Together, these observations raise the possibility of coordinated synaptic remodeling or compensation and identify specific molecular targets for protein-level and functional studies. Relating these targets to phenotype is the next logical step.

Prior behavioral studies provide a useful phenotypic framework for that next step. Earlier work reported no significant novel object location deficit^20^ and identified altered locomotor timing without changes in conventional anxiety-like or aggression measures.^21^ Habituation engages multiple neural systems,^32,33^ and combining transcriptomic profiling with within-animal behavioral measurements in future cohorts could directly connect the present molecular signatures to function. Region-resolved analyses would also enable closer comparison with the earlier dentate-gyrus microglial findings.^20^ In parallel, developmental IL-17A effects^2,4,34^ provide a strong rationale for age-resolved studies that distinguish developmental influences from adult signaling. This broader systems perspective is particularly relevant when considering the complement-related signal centered on *C4b*.

The significant increase in *C4b* provides a specific molecular entry point for examining complement-related mechanisms in this model. Although neither the GO complement activation set nor the Hallmark complement set reached FDR q < 0.05, the GO leading edge contained several complement components, placing the *C4b* result within a broader pathway context. Experimental work in another disease setting links type I interferon responses to neuroinflammation and synapse loss,^35^ and NPTX2–complement interactions provide an additional mechanistic connection.^31^ Together, these observations nominate an immune–complement–synapse axis as a focused, testable hypothesis for this line. Direct measurements of complement proteins, synapse density and microglial engulfment can now evaluate this possibility, while also addressing the main limitations of the current discovery dataset.

The current study therefore defines a clear roadmap for validation. Replication in larger independent cohorts will establish the robustness of the gene-level and gene-set signals identified here, particularly for low-abundance transcripts. Expanding the design to females, additional ages, hippocampal subregions and cell-resolved approaches will determine how broadly these signatures are expressed and will help localize the increased *Rorc* signal and IL-17 receptor expression to specific cell populations. Contemporaneous measurements of circulating IL-17A and other cytokines will strengthen links between peripheral immune state and hippocampal transcription, building on elevations reported in previous cohorts at the same age.^19^ Finally, orthogonal protein assays and targeted perturbations can test the prioritized interferon, UPR, synaptic and complement mechanisms. These next steps build directly on the present transcriptomic resource and provide a practical path from association to mechanism.

In conclusion, hippocampal RNA sequencing in RORγt-transgenic mice revealed coordinated interferon-response and UPR-associated transcriptional signatures together with preferential WT-side enrichment of selected synaptic and calcineurin-related gene sets and FDR-significant candidates including *C4b, Xbp1, Nptx2* and *Mt2*. By integrating gene-level and gene-set analyses, this study provides a focused molecular framework for investigating how sustained Th17 bias may intersect with hippocampal immune, proteostatic and synaptic programs. The dataset is deliberately hypothesis-generating and prioritizes specific cellular, biochemical and functional experiments for the next stage of investigation.

## ABBREVIATIONS

CNS: central nervous system
EDGE: Empirical Analysis of DGE tool in CLC Genomics Workbench
ER: endoplasmic reticulum
ES: enrichment score
FC: fold change
FDR: false discovery rate
FWER: family-wise error rate
GO: Gene Ontology
GO-BP: Gene Ontology biological process
GSEA: gene set enrichment analysis
IFN: interferon
IL-17A: interleukin-17A
LE: leading edge
MSigDB: Molecular Signatures Database
NES: normalized enrichment score
PCA: principal component analysis
poly(I:C): polyinosinic–polycytidylic acid
QC: quality control
RIN: RNA integrity number
RNA-seq: RNA sequencing
RORγt: retinoic acid-related orphan receptor γt
RPKM: reads per kilobase of transcript per million mapped reads
rRNA: ribosomal RNA
SD: standard deviation
Tg: RORγt transgenic
Th17: T helper 17
TSV: tab-separated values
UPR: unfolded protein response
WT: wild type

## ACKNOWLEDGMENTS

The authors thank Kyoko Kishi and Masae Ohtsuka for their technical support. The authors also thank Kentaro Itagaki, Sara Kamiya, Yui Kurama for valuable discussions and critical comments on this study. The authors also acknowledge the Organization for Open Facility Initiatives, University of Tsukuba, for providing access to shared research facilities.

## AUTHOR CONTRIBUTIONS

TS: Conceptualization, Methodology, Validation, Formal analysis, Investigation, Resources, Data curation, Writing – original draft, Writing – review & editing, Visualization, Supervision, Project administration, and Funding acquisition. KN: Methodology, Validation, Formal analysis, Investigation, Data curation, Visualization, and Writing – review & editing. SS: Validation, Investigation, Data curation, and Writing – review & editing. MM: Resources and Writing – review & editing. SI: Resources and Writing – review & editing. YT: Conceptualization, Resources, Supervision, Project administration, and Writing – review & editing. All authors read and approved the final manuscript.

## ETHICS STATEMENT

Animal procedures were conducted under University of Tsukuba institutional oversight and in accordance with institutional animal-care guidelines and the NIH Guide for the Care and Use of Laboratory Animals, as described in Materials and Methods.

## DATA AND CODE AVAILABILITY

Processed gene-level expression and statistical results, sample metadata, complete exported GSEA results and leading-edge memberships are provided as Supplementary Tables S1–S4 and accompanying source-data files. Deposition of the raw sequencing reads in GEO/SRA is in progress; the accession will be added in a revised preprint once available. Analysis scripts and exact archived execution files are being organized for public release; reconstructed exports are distinguished from archived run outputs.

