## Supplementary Materials for "Hippocampal transcriptomic profiling reveals interferon, unfolded protein response, and synaptic signatures in RORγt-transgenic mice"

*Figures S1-S3 and legends for Tables S1-S4*

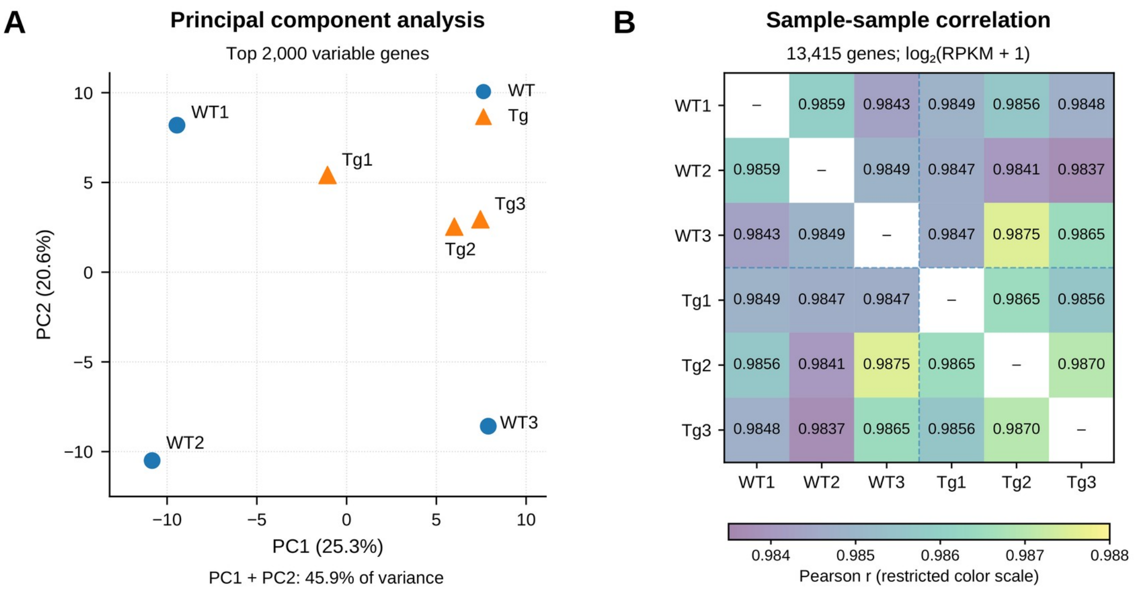


**Figure S1. Principal component analysis and sample–sample correlations of hippocampal RNA-seq expression profiles.**

WT1–WT3 and Tg1–Tg3 represent individual mice (n = 3 per genotype). Genes with mean RPKM > 0.5 across all six samples were retained (13,415 genes) and transformed as $\log_2$(RPKM + 1). (A) PCA of the 2,000 genes with the highest variance in transformed expression. Gene values were centered across samples without unit-variance scaling. Circles denote WT and triangles denote Tg. PC1 and PC2 explain 25.3% and 20.6% of variance, respectively (45.9% combined). (B) Pearson correlations calculated across all 13,415 QC-filtered genes. Values are shown to four decimal places; the 15 distinct pairs range from 0.9837 to 0.9875. Self-correlations are masked and marked with a dash. The color scale is restricted to 0.9835–0.9880 to resolve small differences. All six samples are retained; Rorc is not excluded from the QC-filtered gene set, in contrast to the 13,813-gene GSEA input. No inferential test of within-genotype versus between-genotype correlations was performed. Abbreviations: PCA, principal component analysis; QC, quality control; RPKM, reads per kilobase of transcript per million mapped reads; WT, wild type; Tg, RORγt transgenic.

**Figure S2** **All significant GO biological processes**

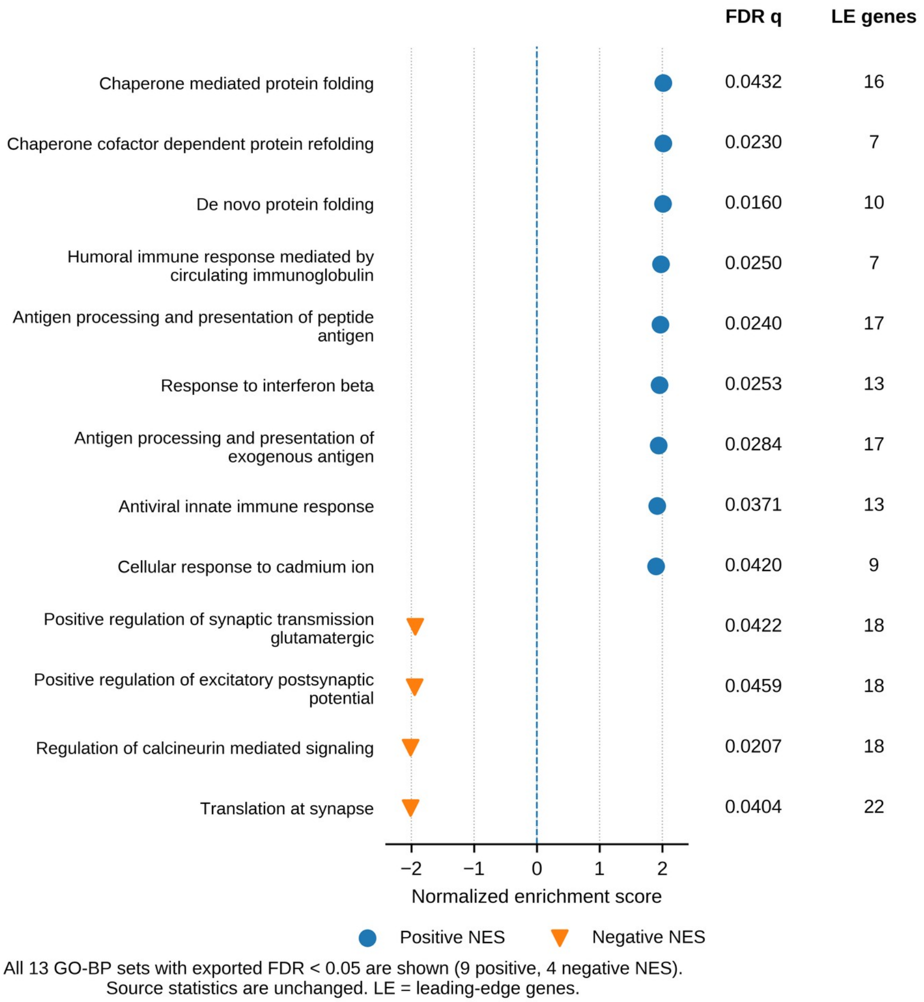


**Figure S2. All significant GO biological-process gene sets.**

All 13 GO-BP terms meeting exported FDR q < 0.05 are shown: nine have positive and four have negative NES. Circles and downward triangles distinguish the two directions. The horizontal position indicates NES; the adjacent columns give the original q value and leading-edge (LE) count. All 13 sets have at least five LE genes. The plot includes cellular response to cadmium ion, which is not shown among the representative terms in Figure 3A. This term name does not demonstrate cadmium exposure. All 3,595 GO-BP results are available in Table S3; no q values were recalculated for the displayed subset. Abbreviations: GO-BP, Gene Ontology biological process; FDR, false discovery rate; NES, normalized enrichment score; LE, leading edge.

**Figure S3** **Overlap among enriched gene sets**

3 Hallmark and 13 GO-BP sets with exported FDR < 0.05

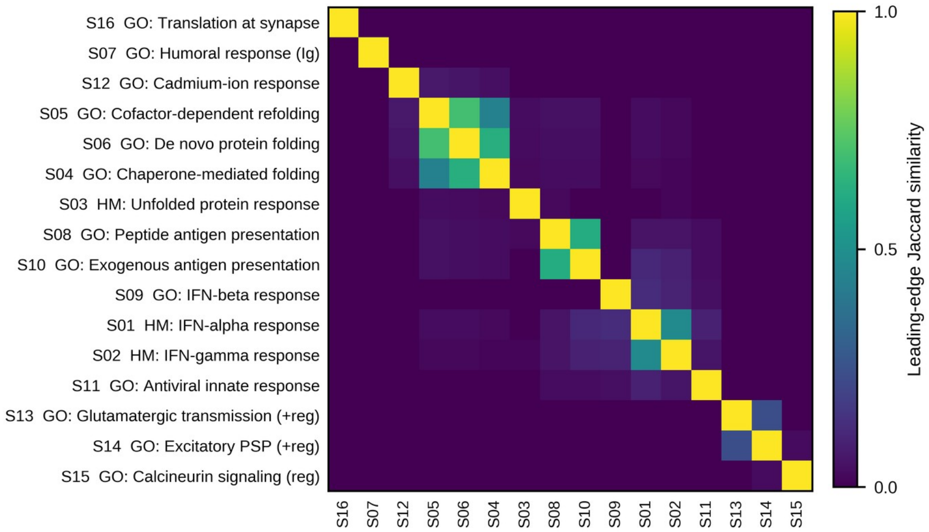


Average-linkage ordering of 1 − Jaccard. Full term names are provided in Table S4.
This is leading-edge overlap, not GO semantic similarity or a significance test.

**Figure S3. Leading-edge overlap among significant Hallmark and GO-BP gene sets.**

Jaccard similarity, $|A \cap B|/|A \cup B|$, was calculated for all pairs among the three significant Hallmark and 13 significant GO-BP leading-edge lists. The matrix was reordered by average-linkage clustering on 1 − Jaccard, with optimal leaf ordering. Zero indicates no overlap and one indicates identical lists; the diagonal is retained at one. Stable IDs map abbreviated labels to the full terms in Table S4. This is a descriptive analysis of leading-edge membership, not GO semantic similarity or an additional significance test. Shared genes mean that related enrichments are not independent evidence. Significant refers to the original false discovery rate (FDR) $q < 0.05$ within each collection. GO-BP denotes Gene Ontology biological process.

### SUPPLEMENTARY TABLE LEGENDS

**Table S1. Sample metadata, RNA integrity and mapping summary.**

The table retains figure labels, analysis and original QC identifiers, animal IDs, preparation labels, run IDs, RIN values, reported NanoDrop values, RNA input and mapping counts for all six mice. Mapped total is the sum of reads mapped in pairs and in broken pairs. Percentages were calculated from the original integer counts; the source unit is reads, not read pairs. NanoDrop units were not stated and have not been inferred. Sex, age and tissue are taken from the study description. Original QC preparation labels and run-index labels are retained as separate fields pending crosswalk verification. The differing TSK and TKS prefixes are not silently replaced. Abbreviations: QC, quality control; RIN, RNA integrity number; ID, identifier.

**Table S2. Complete gene-level results and selected subsets.**

The full TSV contains 49,585 gene rows, individual total counts and RPKM, original EDGE P/FDR values, CLC-signed ratios and weighted differences, and explicitly derived expression summaries. The Excel companion contains the 16 genes meeting FDR < 0.05 and 212 exploratory candidates meeting unadjusted P < 0.05 with |CLC-signed ratio| ≥ 2. $Log_2$ fold change is sign(FC_CLC) × $log_2$(|FC_CLC|). Genotype SDs use the sample standard deviation across three mice. GSEA-filter and QC-filter flags use, respectively, mean RPKM > 0.5 in either genotype and mean RPKM > 0.5 across all six samples. Original inferential statistics are preserved. These expression-filter flags identify eligible genes before the additional Rorc exclusion used for GSEA; final analysis membership is recorded by used_in_GSEA in Table S3. Abbreviations: EDGE, Empirical Analysis of DGE in CLC Genomics Workbench; FC, fold change; FDR, false discovery rate; GSEA, gene set enrichment analysis; QC, quality control; RPKM, reads per kilobase of transcript per million mapped reads; SD, standard deviation; TSV, tab-separated values.

**Table S3. Complete exported gene set enrichment results.**

All 5,198 exported result rows are retained: 50 Hallmark, 3,595 GO-BP and 1,553 mixed curated sets. The Mixed_curated worksheet is the renamed Reactome_WikiPathways source worksheet and contains 662 REACTOME, 117 WP, 82 BIOCARTA and 692 other sets, including named experimental signatures. ES, NES, nominal P, FDR q, FWER P, Tag, Gene percentage, nLE and complete leading-edge strings are preserved. Original q values retain their collection-specific context and are not recomputed by term prefix. Exported zero nominal P values denote permutation estimates. The Ranked_gene_list worksheet retains all 13,814 expression-filtered genes before Rorc exclusion; used_in_GSEA is TRUE for the 13,813 included genes and FALSE for Rorc. The GSEA_input_noRorc worksheet and accompanying two-column RNK export contain only the TRUE rows, in descending signed-score order. Abbreviations: ES, enrichment score; NES, normalized enrichment score; FDR, false discovery rate; FWER, family-wise error rate; GO-BP, Gene Ontology biological process; nLE, number of leading-edge genes. Tag and Gene percentage retain their source output labels.

**Table S4. Leading-edge memberships, gene expression and set overlap.**

For the 26 sets meeting FDR q < 0.05 across the three exported collections, 493 gene–set memberships represent 280 unique genes. Exact source gene symbols link memberships to gene-level EDGE statistics and individual RPKM values. Set-level significance does not imply significant differential expression of every member. The workbook includes the 16-set key, 120 distinct pairwise comparisons and Jaccard matrix for Figure S3. The compressed TSV contains all 96,884 one-gene-per-row leading-edge memberships across the full set of 5,198 exported results. Abbreviations: EDGE, Empirical Analysis of DGE in CLC Genomics Workbench; FDR, false discovery rate; RPKM, reads per kilobase of transcript per million mapped reads; TSV, tab-separated values.
